# Combining modelling and experimental approaches to assess the feasibility of developing rice-oil palm agroforestry system

**DOI:** 10.1101/2022.12.19.521043

**Authors:** Raphaël P.A Perez, Rémi Vezy, Romain Bordon, Thomas Laisné, Sandrine Roques, Maria-Camila Rebolledo, Lauriane Rouan, Denis Fabre, Olivier Gibert, Marcel De Raissac

## Abstract

Climatic hazards affecting the main rice producing regions of Indonesia increase the risk of annual production loss and encourage the development of innovative strategies to maintain stable production. Conversion of oil palm monocultures to rice-based intercropping systems is a strategy to be considered, but relies on the existence of suitable planting management that optimizes both palm productivity while providing enough light for undergrowth rice varieties tolerant to shady conditions. This paper proposes to couple a model of light interception on virtual canopies with indoor experiments to evaluate the feasibility of developing rice-oil palm agroforestry systems. We first selected a planting design that optimized the transmitted light available for rice using a functional-structural plant model (FSPM) of oil palm. Secondly, we reproduced the light regime simulated with specific changes in the intensity and the daily fluctuation of light in controlled conditions. Three light treatments were designed to test independently the effect of daily light quantity and the effect of diurnal fluctuation on contrasted rice subpopulations.

Light quantity was the main factor driving changes in plant morphology and architecture, while light fluctuation only appeared to explain variations in yield components and phenology. This study highlighted the importance of light fluctuation in the grain filling process and resource reallocation. The conservation of relative change among varieties between treatments suggests that varietal responses to low light are likely to be heritable, and that varietal screening under full light can provide clue on varietal behavior under low light. However, the identification of specific traits such as a limited expansion of leaf area and a conservation of leaf senescence under shade and high light fluctuation paves the way for selecting varieties dedicated to agroforestry systems. Further investigations including light quality and larger genotypic population to screen are discussed.

## Introduction

The recurrent droughts caused by the ‘El Niño’ phenomenon have a significant impact on both rice and oil palm cultivation in Southeast Asia, particularly in Indonesia. These climatic hazards affecting the main rice-producing regions (Java and Bali), increase the risk of annual production loss (Naylor *et al*. 2007) and encourage the development of innovative strategies to maintain stable production. Oil palm yields also decrease in regions where more and more drought periods are recorded (South Sumatra), and thus raise interest in limiting competition for water through adapted planting density. Subsequently, oil palm/rice intercropping in the rainy season alternating with palms alone at reduced density in the dry season could be a strategy to extend suitable area for rice while reducing competition for water between palms during dry spells. The feasibility of such a system depends on a specific management of the resources depending on the two species and seasons: the water availability for oil palm during the dry season and the transmitted light for rice during the rainy season. A main concern is therefore on adjusting the palm density to maintain high yields of both crops, and on screening rice varieties best adapted to shady conditions.

Research on oil palm agroforestry systems is rapidly expanding in the scientific literature (Bhagwat & Willis 2008; Khasanah *et al*. 2020). The majority of these studies focused on the ecological impact of these systems compared to conventional monoculture systems (Gérard *et al*. 2017; Ashraf *et al*. 2018; Zemp *et al*. 2019; Koussihouèdé *et al*. 2020; da Silva Maia *et al*. 2021). Rare studies highlighted the interest of palm agroforestry systems for their good agronomic performance and economic profitability on the one hand (Gawankar *et al*. 2018; Ahirwal *et al*. 2021), but also for buffering extreme climatic events and thus improving the sustainability of these systems (Ashraf *et al*. 2019). However, very few studies proposed intercropping with rainfed rice (Alridiwirsah *et al*. 2019), a crop that is nevertheless predominant in Asia. Oil palm-based intercropping is often considered at juvenile stage of oil palm, as a transitory option for a couple of years, since less than five percent of the incident light forages the canopy when oil palms reach maturity at a conventional density of 136 or 143 plants ha^-1^. So far, no study investigated how changes in planting patterns and density of oil palm could be optimized for intercropping purposes, because i) such study requires years and strong financial support, and ii) suitable planting management that optimizes palm productivity while providing enough light for undergrowth crop is not straightforward. The recent development of modelling tools, using 3D models of plants in combination with light modelling (Perez *et al*. 2022) now opens the way to test how planting design would modulate light availability under oil palm canopy without involving expensive trials.

Despite the growing interest in rice-based agroforestry systems (Wangpakapattanawong *et al*. 2017; Rodenburg *et al*. 2022; Masure *et al*. 2022), the diversity of rice response to fluctuations in radiation as well as low light remains unexploited. Up to now, varietal improvement programs in rice have focused on the selection of productive materials in full sun conditions. The existence of genetic variability in response to photosynthesis induction rate during shade/light alternations (Qu *et al*. 2016; Acevedo-Siaca *et al*. 2020; Taniyoshi *et al*. 2020), and independently to shade tolerance (Wang *et al*. 2015a; Dutta *et al*. 2018), suggest that it could be possible to select rice material suitable for agroforestry. Nevertheless, these studies investigated shade tolerance using continuous artificial light attenuation with shade nets, often starting at the reproductive stage of plant growth, thus preventing mimicking light conditions in agroforestry, where undercrop can be exposed to full sunlight around noon. In these systems, the light environment is much more complex than a constant light attenuation, since both effects of shading and daily light fluctuation can affect plant growth, from germination to harvesting.

This study raises the question of whether rice genotypes selected to be most efficient under optimal radiation are most efficient when the amount and the dynamics of radiation are altered, as found in agroforestry systems. Given the nonlinear relationship between light interception and photosynthesis, not only the reduction in light quantity but also the changes in daily light fluctuation can affect resource acquisition and subsequently plant growth, development and yield. Using a functional-structural plant model (FSPM) estimating light transmission through 3D canopy, we first simulated how palm density and planting design affect the quantity of transmitted light to rice, and selected a pattern optimizing light foraging for rice and oil palm density. Secondly, we reproduced in controlled conditions the specific changes in light intensity and fluctuation obtained in the simulation. Three light treatments differing in either the quantity or the dynamics of radiation were thus set up to investigate changes in phenological, morphological, physiological, and productive traits including physicochemical traits on eight contrasted rice varieties.

## Material and Method

### Modelling light transmission in innovative oil palm-based agroforestry system

We can test how planting design can modulate light availability under oil palm canopy by coupling a 3D structural model of oil palm (Perez *et al*. 2016, 2018) with a light interception model (Dauzat & Eroy 1997). This modelling approach helps to compute the light transmitted over time and space according to any design and thus allows the exploration of innovative and suitable planting patterns for agroforestry systems. In this study, we assumed that the architectural plasticity of oil palm had negligible impact on light capture in comparison to the changes in light interception due to planting patterns, as concluded in a previous study (Perez *et al*. 2022). A simulation study was performed with 3D mock-ups of oil palm representing plant architecture at the mature stage (8 years after planting), and displayed in rows defined by two parameters: the distance between rows and the distance between plants within rows (Fig S1). Eight planting patterns were designed *in silico* to estimate the light transmitted through the palms in a horizontal plane placed one meter above the ground (Table 1).

**Table 1:** The eight designs tested with the corresponding distance between plants and density. The two parameters to define a design are the distance between rows (dist inter) and within rows (dist intra).

| dist intra (m) | dist inter (m) | density (plants ha<sup>-1</sup>) |
| --- | --- | --- |
| 8 | 14 | 89 |
| 7 | 14 | 102 |
| 8 | 12 | 104 |
| 7 | 12 | 119 |
| 8 | 10 | 125 |
| 8 | 9 | 138 |
| 7 | 10 | 142 |
| 7 | 9 | 158 |

Distances within rows (7 and 8 m) were selected according to the average crown dimensions of mature oil palm, with the objective to mutual shading. Distance between rows (9, 10, 12, 14m) were selected to modulate planting density from low (89 plants ha^-1^) to high density (158 plants ha^-1^). Simulations were done using radiative data collected in South Sumatra on March 20th, 2019 (sunny day), with a global radiation reaching 28.4 MJ m^-2^. The light transmitted under the canopy was integrated over the day for every element of the horizontal plane (each element is a 16 cm x 16 cm square). Considering the space for growing rice at least 3 m distant from oil palm rows, we estimated on average photosynthetically active radiation (PAR) available in the rice area over a day.

### Reproducing the simulated agroforestry light conditions in growth chambers

The experiment was carried out in the Abiophen platform of Montpellier in controlled conditions (Aralab Fitoclima 25000HP Growth chamber) in which light intensity and time course were adjusted while maintaining equivalent temperature and air humidity (Fig S2). Three light treatments (light supplied using Philips Master Colour CDM-T Elite MW 315W) were designed to test independently the effect of daily light quantity and the effect of diurnal fluctuations of light on rice. The three light treatments were set up in three independent growth chambers, respectively:

- a control treatment (C) following the dynamic of incident light observed in Sumatra (with a ceiling radiation value that can be reached *i.e* 1100 μmol_photons_ m^-2^ s^-1^)
- the agroforestry shading regime corresponding to the planting design selected from the simulation study (S-AF)
- a uniform shading treatment (S)

The light reduction in the S treatment was performed to obtain the same amount of total daily radiation than in the S-AF treatment (Fig 1A), and corresponded to a constant shade that could be obtained with a shading net. We characterized this amount of light by integrating the photosynthetic photon flux density (PPFD) over a day, hereafter called daily light integral (DLI, Poorter et al. 2019). The DLI were 32.4 mol m^-2^ d^-1^, 12.9 mol m^-2^ d^-1^ and 12.9 mol m^-2^ d^-1^ for C, S-AF and S, respectively, both shading treatments thus providing equivalently 60% of light attenuation (Fig 1B). Regarding light fluctuation, we calculated an index of daily light fluctuation (DLF) as the daily difference between the maximal and the minimal PPFD (i.e 0 μmol.m^-2^ s^-1^). DLF for S and S-AF were 478 and 694 μmol m^-2^ s^-1^, thus representing respectively a reduction of 52% and 30% of light fluctuation compare to the control treatment. Light spectra were also compared in the three growth chambers to ensure similar light quality among treatments (Fig S3).

**Figure 1:**
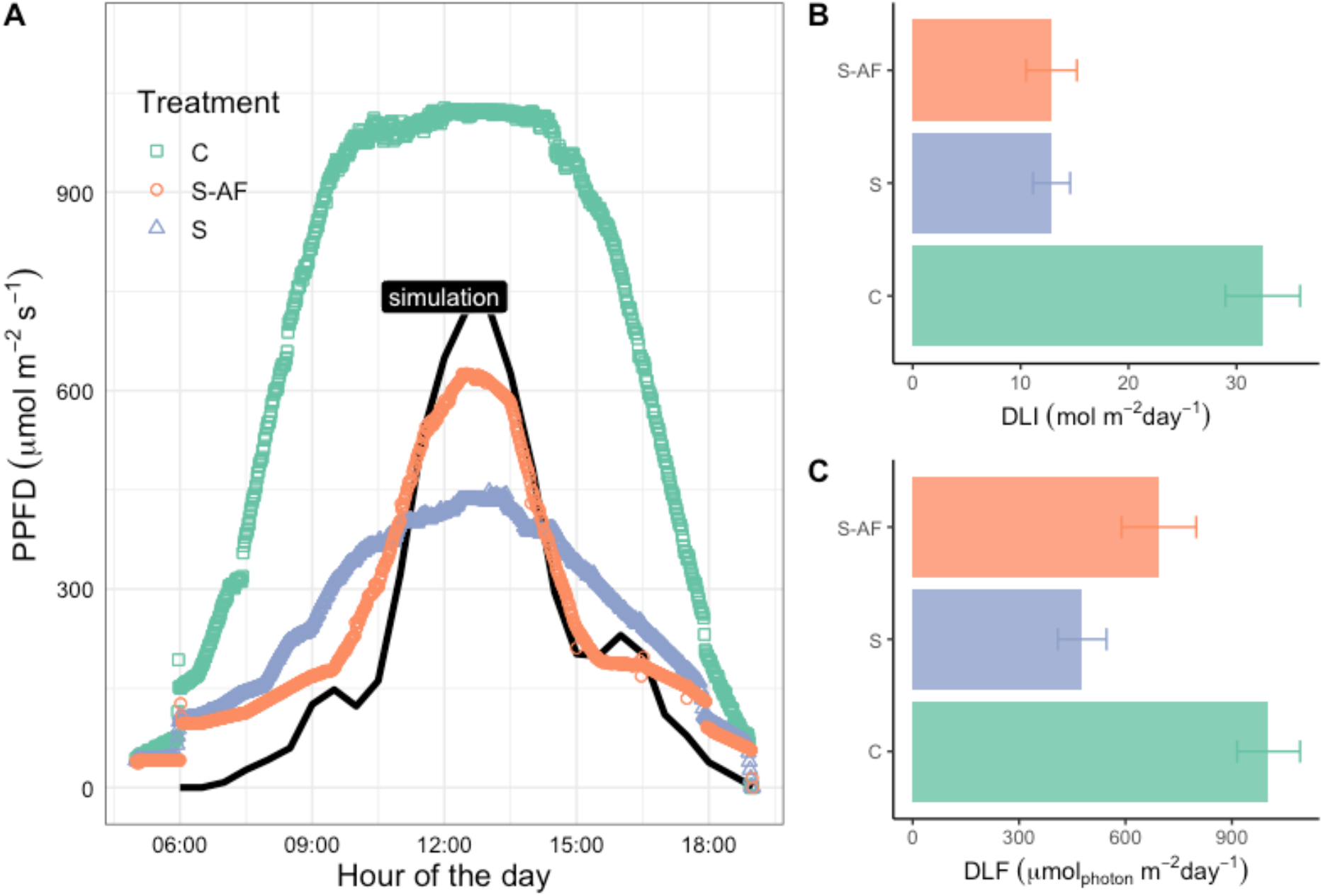
A) Mean diurnal time course of the photosynthetic photon flux density (PPFD) measured under three light treatments in the phytotrons during the whole cycle of plant growth (C: control. S-AF: agroforestry-like shade. S: constant shade). The black line indicates the PPFD simulated in silico in the 7m x 12m planting design. B) Daily light integral (mean and inter-days standard deviation) measured for the three treatments. S and S-AF represent 60% loss of radiations in comparison to C. C) Daily light fluctuation (mean and inter-days standard deviation) measured for the three treatments. S and S-AF represent respectively 52% and 30% loss of light intensity in comparison to C.

### Plants material and phenotypic monitoring

Eight accessions of rice representing diversified subpopulations and geographical regions were provided by the seeds and genetic resources laboratory (TBRC) held in Montpellier, France (Table 2). Seeds were sown in individual pots of 4.8L (16cm x 16cm x 23cm) and fertilized using 3g L^-1^ of NPK fertilizer (50% of Basacote 13-5-18 and 50% of Bio 5-3-8; Compo – expert). In each of the three growth chambers, accessions were grouped in tables of 28 plants (corresponding to a density of 32 plants m^-2^) to prevent any competition for light among varieties within the table. Eight tables were placed in each phytotron, and pots were kept flooded to prevent any water stress. Temperature and relative humidity followed the average value observed in South Sumatra: temperature varied from 22°C by night up to 32°C during the day and relative humidity from 83% down to 65% during the day (Fig S2). Temperature and vapour pressure deficit were monitored similarly in the three treatments so that phenotypic variations were interpreted as the consequence of light treatment alone.

**Table 2:** Rice accessions monitored during the experiment in controlled conditions. *accessions with full phenotypic monitoring from sowing to harvesting.

| Accession | Ref | Gervex | Subpopulation | Origin |
| --- | --- | --- | --- | --- |
| JC1 | V1 | 9 091 | Basmati | India |
| MALAGKIT PIRURUTONG | V6 | 8 182 | Japonica | Philippines |
| IR64 * | V3 | 8 531 | Indica | Philippines |
| TREMBESE * | V7 | 43 675 | Japonica | Indonesia |
| SINNA SITHIRA KALI | V4 | 51 064 | indica | Sri Lanka |
| IRAT 177 * | V8 | 7 415 | Javanica | Guyane |
| BRASILEIRO * | V5 | 809 | Japonica | Brasil |
| CASAMANCE V2 | V2 | 1123 | Glaberrima | Sénégal |

Phenotypic monitoring was conducted from sowing to harvesting, with phenological, morphological, physiological and yield measurements (Table 3). On each table, the 10 inner plants among the 28 plants constituting the canopy were studied to prevent border effects in the analyses. Phenological observations (leaf tip appearance on main stem and tillers count) were made once a week. For leaf appearance a Haun scale was used: the leaf was counted once its tip was visible and the total number of emerged leaves was expressed on a decimal scale, with partially emerged leaves counted according to the visible fraction of the full size (Haun 1973; Tivet *et al*. 2001).

**Table 3:**
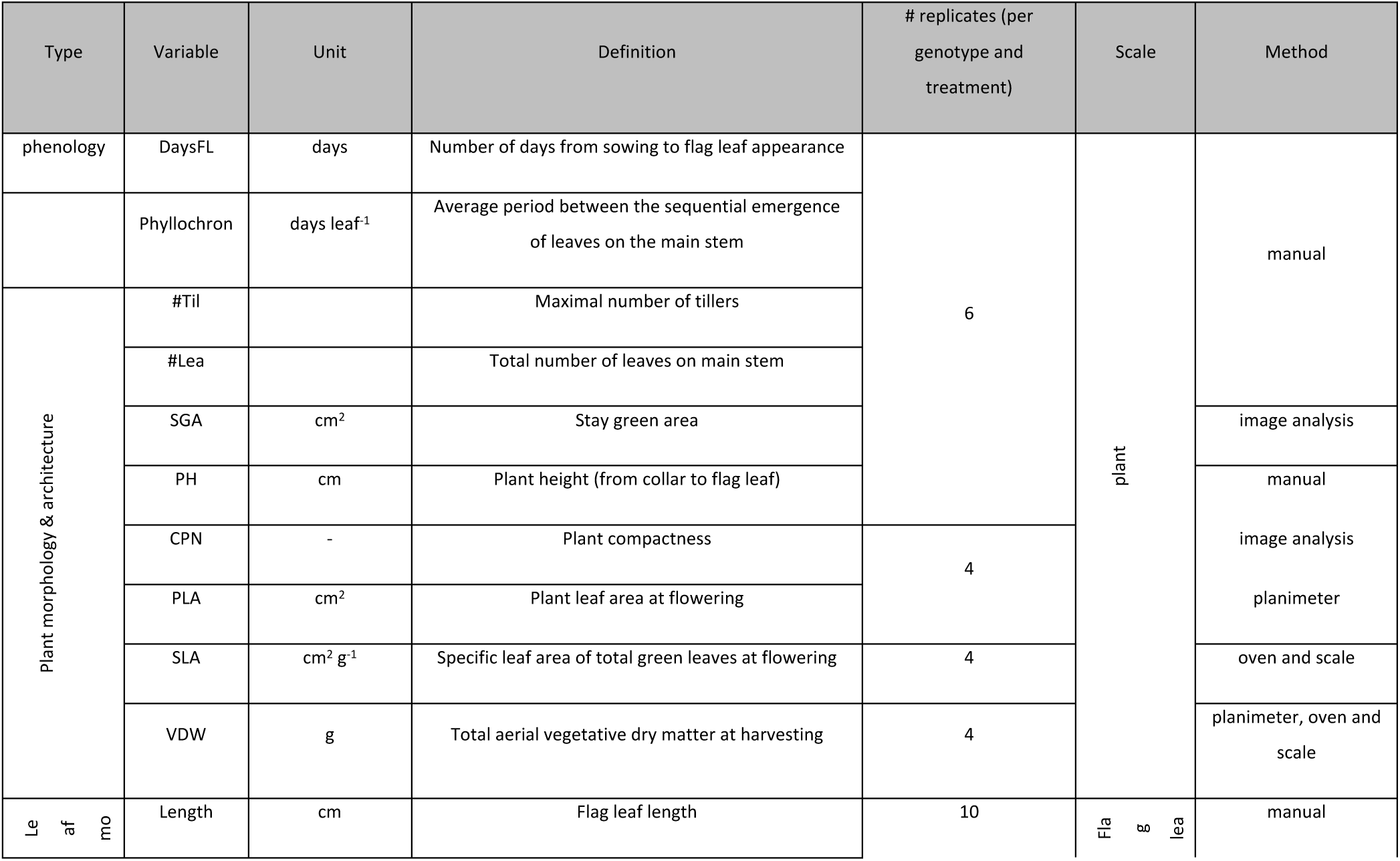
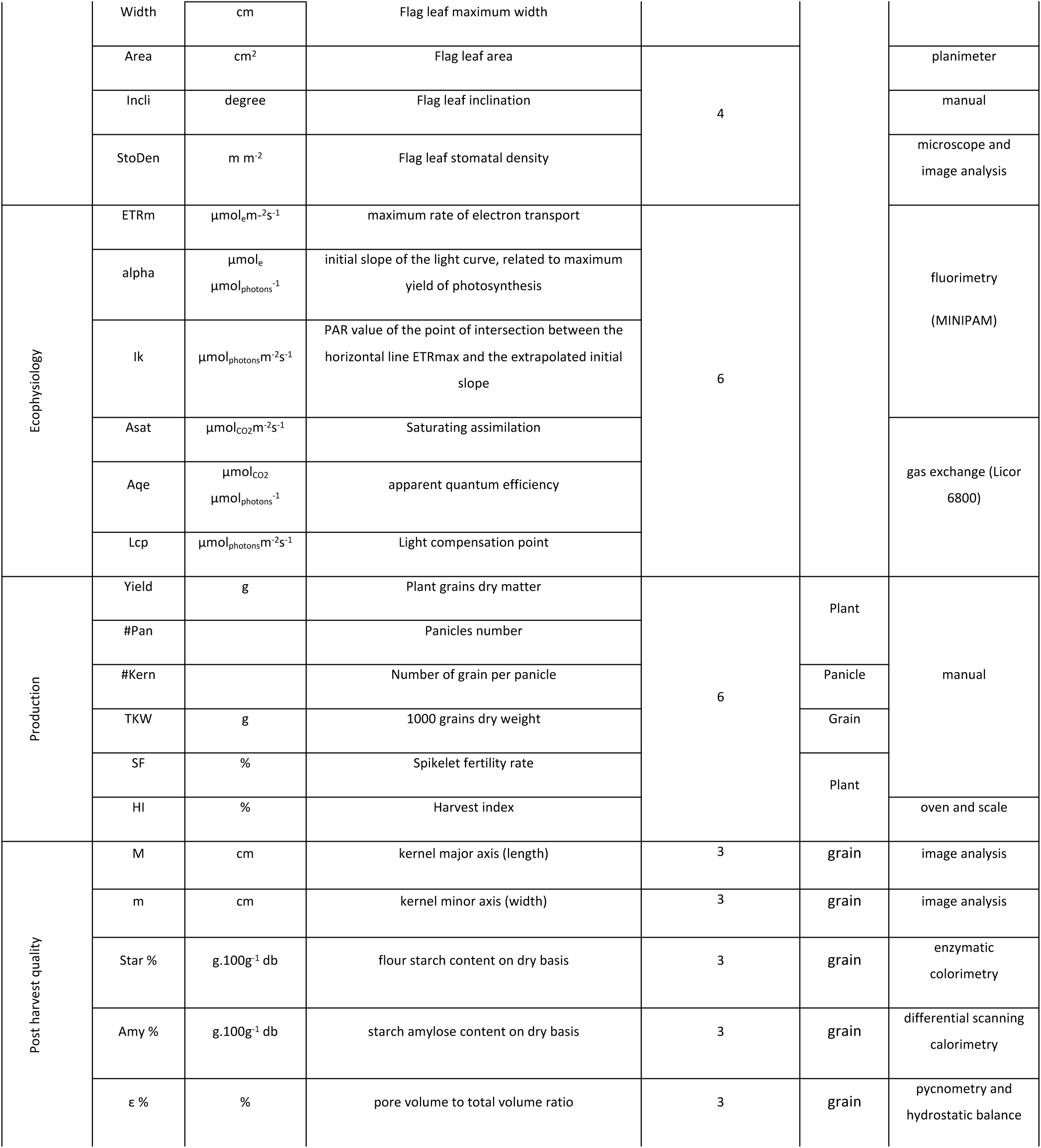
Phenotypic variables collected during the experiment.

**Table 3:** Summary of models analysis to assess the effect of variety, daily light integral (DLI), daily light fluctuation (DLF) and their interaction on phenotypic traits. An ANCOVA is performed with variety as factor and DLI and DLF as covariables (ns: non-significant, *P<0.05;**P<0.01;***P<0.001). For variable abbreviations, see Table 2.

|  | Rep | var | Variety | DLI | DLF | V x DLI | V x DLF | r2 | LRT |
| --- | --- | --- | --- | --- | --- | --- | --- | --- | --- |
| Yield components | 6 | Yield | *** | *** | ns | ns | * | 0.76 | *** |
|  | 6 | #Pan | *** | *** | ns | ns | ns | 0.83 | ns |
|  | 6 | #Kern | *** | ns | ** | ns | * | 0.65 | *** |
|  | 6 | TKW | *** | *** | *** | ** | ** | 0.90 | *** |
|  | 6 | SF | *** | * | *** | ns | * | 0.42 | *** |
|  | 6 | HI | *** | *** | *** | ** | *** | 0.92 | *** |
| Flag leaf morphology | 10 | Length | *** | *** | ns | ns | ns | 0.52 | ns |
|  | 10 | Width | *** | *** | ns | ** | ns | 0.77 | ns |
|  | 10 | LWratio | *** | ns | ns | * | * | 0.56 | ns |
|  | 10 | Area | *** | *** | ns | ns | ns | 0.56 | ns |
|  | 4 | Incli | *** | *** | ns | *** | *** | 0.94 | *** |
|  | 4 | StoDen | *** | *** | ns | ns | ns | 0.73 | ns |
| Phenology | 10 | DaysFL | *** | *** | * | ns | ns | 0.66 | ns |
|  | 10 | Phyllochron | *** | *** | * | *** | * | 0.65 | ** |
| Plant architecture | 10 | #Lea | *** | * | ns | *** | *** | 0.77 | *** |
|  | 10 | #Til | *** | *** | *** | ** | ns | 0.94 | *** |
|  | 4 | PLA | *** | *** | ns | ns | ns | 0.47 | ns |
|  | 4 | CPN | *** | *** | ns | * | ns | 0.84 | ns |
|  | 4 | SLA | *** | *** | ns | ns | ns | 0.64 | ns |
|  | 6 | VDW | *** | *** | * | ns | ** | 0.54 | *** |
|  | 6 | PH | *** | *** | ns | ns | ns | 0.85 | ns |
| Ecophysiology | 6 | Sgi | *** | *** | *** | *** | ** | 0.90 | *** |
|  | 6 | Lcp | *** | ** | * | * | *** | 0.61 | *** |
|  | 6 | Asat | * | ns | ** | ns | *** | 0.34 | *** |
|  | 6 | Aqe | *** | ns | ns | ns | *** | 0.48 | *** |
|  | 6 | Rd | * | ns | * | ns | ** | 0.27 | *** |
|  | 6 | ETRm | ns | ns | * | ns | * | 0.21 | ** |
|  | 6 | alpha | *** | * | ns | ns | *** | 0.49 | *** |
|  | 6 | lk | ** | ns | ** | ns | ns | 0.24 | * |
| Grain morphology | 3 | M | *** | ns | ns | ns | ns | 0.95 | ns |
| & physicochemical quality | 3 | m | *** | ns | ns | ns | ns | 0.48 | ns |
|  | 3 | Star % | * | ** | * | ns | ns | 0.49 | ** |
|  | 3 | Amy % | *** | ns | ns | ns | ns | 0.97 | ns |
|  | 3 | ε % | *** | * | *** | *** | *** | 0.92 | *** |

At flowering, four out of the ten studied plants were collected for image-based analysis and destructive measurements. Destructive measurements consisted in total plant leaf area (PLA) using a planimeter and the shoot vegetative dry matter (leaves and stems). The ratio between PLA and the leaves dry matter gave an estimate of plant specific leaf area (SLA). The stomatal density (StoDen) was observed on flag leaves, from a footprint taken by a nail polish on the leaf surface and counted with ImageJ on the images observed on the microscope (Fig 2A). An index of plant compactness (CPN) was derived from plant image, expressed as the ratio of the projected area of leaves over the projection of the volume defined by the whole aerial part of the plant (Fig 2B).

**Figure 2:**
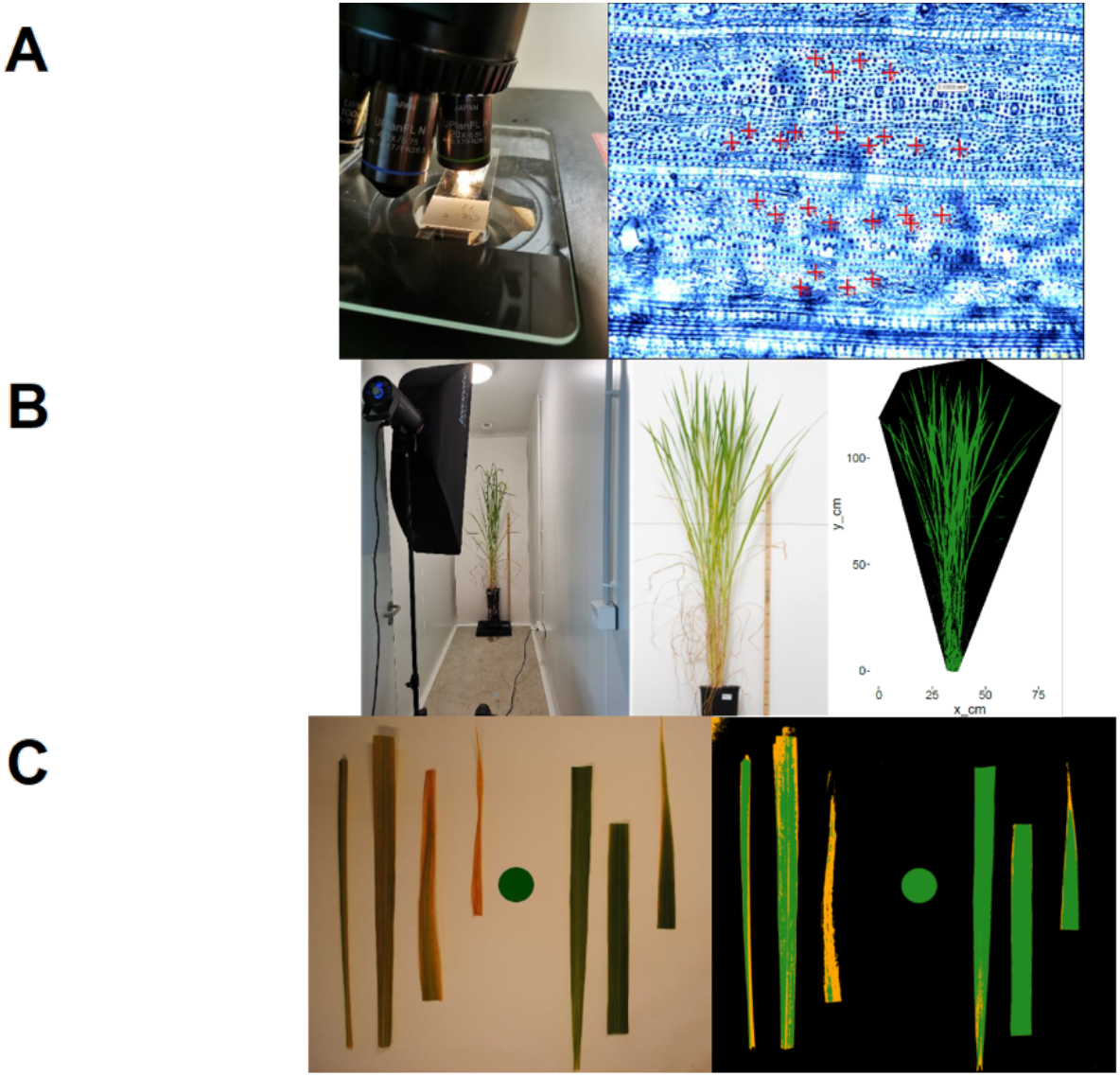
Image-based phenotypic variables. A) Right: Stomatal density observed on microscope slide, left: microscope image with stomata indicated with red crosses within a fixed area of 0.1 mm^2^. B) Whole plant picture to retrieve compactness. Compactness is estimated as the ratio between the number of green pixels over the total pixels defining the convex hull of the areal part of the plant (green and black pixels). C) Estimation of the stay green area (SGA) from a picture of green leaves at harvesting. SGA corresponds to the area in green defined by green pixel. The green coin in the middle is used as a green reference for the segmentation process and scaling the pixel.

At harvesting, the six remaining plants were dissected into vegetative parts and reproductive parts. On the vegetative part, the total dry mass was weighted (vegetative dry weight: VDW) and pictures of the green leaves were used to estimate a stay green area (SGA) from pixel segmentation, defined as the green area left on the main stem (Fig2 C). On the reproductive part, the yield was measured as the total grains dry matter collected on all the panicle of the plant. The panicle of the main stem was used to count the number of fertile and unfertile grain per panicle and derive a spikelet fertility rate (SF). The harvest index (HI) was estimated as the ratio of the yield over the total aerial biomass (vegetative and panicles structural components).

Further analyses were performed on grain morphological and physicochemical feature on a subsample of the harvested spikelets. Paddy kernels were oven-dried at 45°C in a ventilated oven until constant weight was reached. Kernels were then dehusked and polished using a Satake model laboratory testing husker THU35C-T and Tadd mill 4E-230 machines. Digital RGB images (800 dpi) of polished kernels were acquired using a flatbed scanner (Epson Perfection v700 Photo). The grain major (length *M)* and minor axis (width *m)* were computed from the projected area of the binarized image of individual kernels fitted from an ellipse, using ImageJ analysis software platform. Fine-ground rice flours were obtained using cargo kernels a ball mill (Dangoumill 300, Prolabo, France). After fine-group flour moisture content determination by thermogravimetry at 104°C, total starch content (%db) was estimated in triplicate by enzymatic colorimetry (515 *n*m) from the total released glucose after incubation and hydrolysis of about 300 mg db of flour with thermostable-α-amylase (Novo Nordisk, Denmark), and later with glucose oxidase and peroxidase (GOD & POD, Sigma, USA), as per Giraldo Toro et al. (2015). The trace of free glucose was neglected in the native rice flours. The amylose content was then estimated in duplicate using Differential Scanning Calorimetry DSC 8500 (Perkin Elmer, USA) as per Pérez et al. (2013) with modifications. The enthalpy change of formation of complexes between amylose and phospholipids during cooling was estimated from 10-11mg db of flour mixed to 40μl of lysophospholipid 2% (w/V in water) against a pure amylose standard. The kernel porosity (ε = 1-*app d/d*) expressed as percentage was then computed based on the estimation of the true density (*d* in g.cm^-3^) of fine-ground flourby pycnometry using helium carrier and the apparent density (app d in g.cm^-3^) using few kernels on a hydrostatic balance with ethanol. and expressed as percentage.

Gas exchange measurements were conducted on the flag leaf few days after flag leaf emission on the six plants monitored until harvesting. Response curves to photosynthetic photon flux density (PPFD) were performed using a portable photosynthesis system (LI-6800; LiCor Biosciences, Lincoln, NE, USA) at around 10 a.m. each morning over a week. Assimilation-irradiance (A – PPFD) curve fitting analysis were then conducted using the Mitscherlich function (Eq 1) (Heschel *et al*. 2004) to estimate the three photosynthetic parameters: the light saturated assimilation (A_sat_), apparent quantum yield (Aqe) and the light compensation point (L_CP_) for each individual.

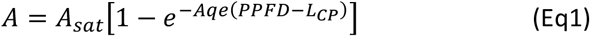

Electron transport rate - irradiance (ETR-PPFD) curve fitting analysis were also determined using equation (Eq 2) to extract the maximum electron transport rate (ETR_m_), the quantum efficiency of photosynthesis (α) and the minimum saturating irradiance (I_k_=ETR_m_/ α).

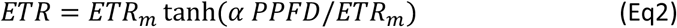

### Assessing the effect of daily light integral (DLI) and daily light fluctuation (DLF) on phenotypic traits

The three treatments considered in this study varied either from the amount of light receive (DLI) and/or in the intensity of light fluctuation perceived over a day (DLF). Each light treatment was a single combination of DLI and DLF. To evaluate if the difference in phenotypic variations were due to DLI or DLF, we conducted statistical analysis based on analysis of covariance models (ANCOVA). For each phenotypic variable studied, we tested the two following ANCOVA models:

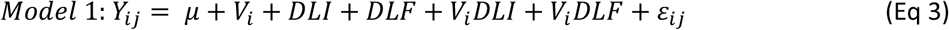

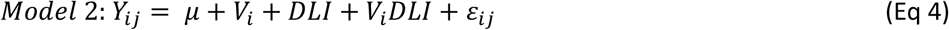

Where Y_ij_ is the phenotypic value of the individual j of the variety i; μ is model grand mean; V_i_ is the effect of the variety I; DLI and DLF are covariates, V_i_DLF and V_i_DLI are the interaction terms and is the residual term. Normality of model residuals and homoscedasticity were tested, and response variables were transformed using square-root or Box-Cox transformation accordingly to ensure normality. For each phenotypic trait, we conducted model testing by comparing the two models fit using a likelihood ratio test (LRT):

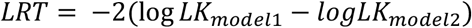

Were logLK is the model log likelihood. LRT was compared with a Chi-squared distribution to obtain a p-value. Significant differences between the two nested models (p-value<0.05) meant that DLI alone did not explained the phenotypic variations observed, and therefore that considering the changes in light fluctuation (DLF) improved model prediction. Likelihood ratio tests were performed using the lrtest function of the lmtest R package (Zeileis & Hothorn 2002).

## Results

### Defining the light regime available for growing rice under an oil palm-based agroforestry system

The simulation of light transmission on the eight planting patterns investigated clearly demonstrated the significant reduction in the light transmitted with increasing planting density (Fig 3). For the lowest density, light transmission reached a maximum of 43% in the middle of the inter row, while it only reached 13% for the highest density. Simulation outputs showed an important decrease in radiation with increasing palm density, with a maximum irradiance around midday. The daily radiation available for rice decreased with planting density (Fig 4), the selection of an interesting design hence depended on the trade-off between oil palm density and daily PAR transmitted for rice.

**Figure 3:**
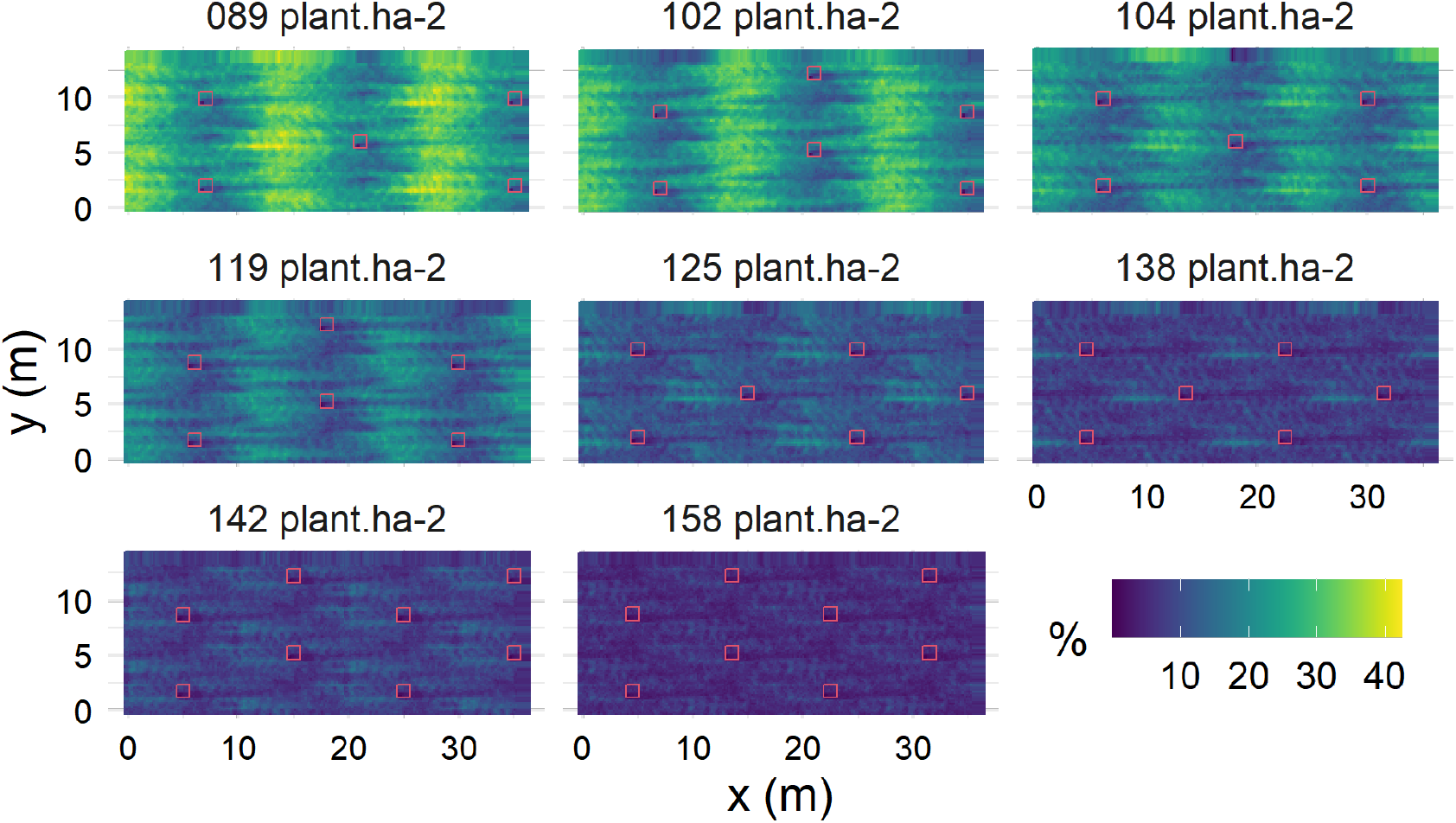
Simulated light transmission under oil palm canopies with various planting designs. Colors indicates percentage of incident light and red squares the positions of palm trees.

**Figure 4:**
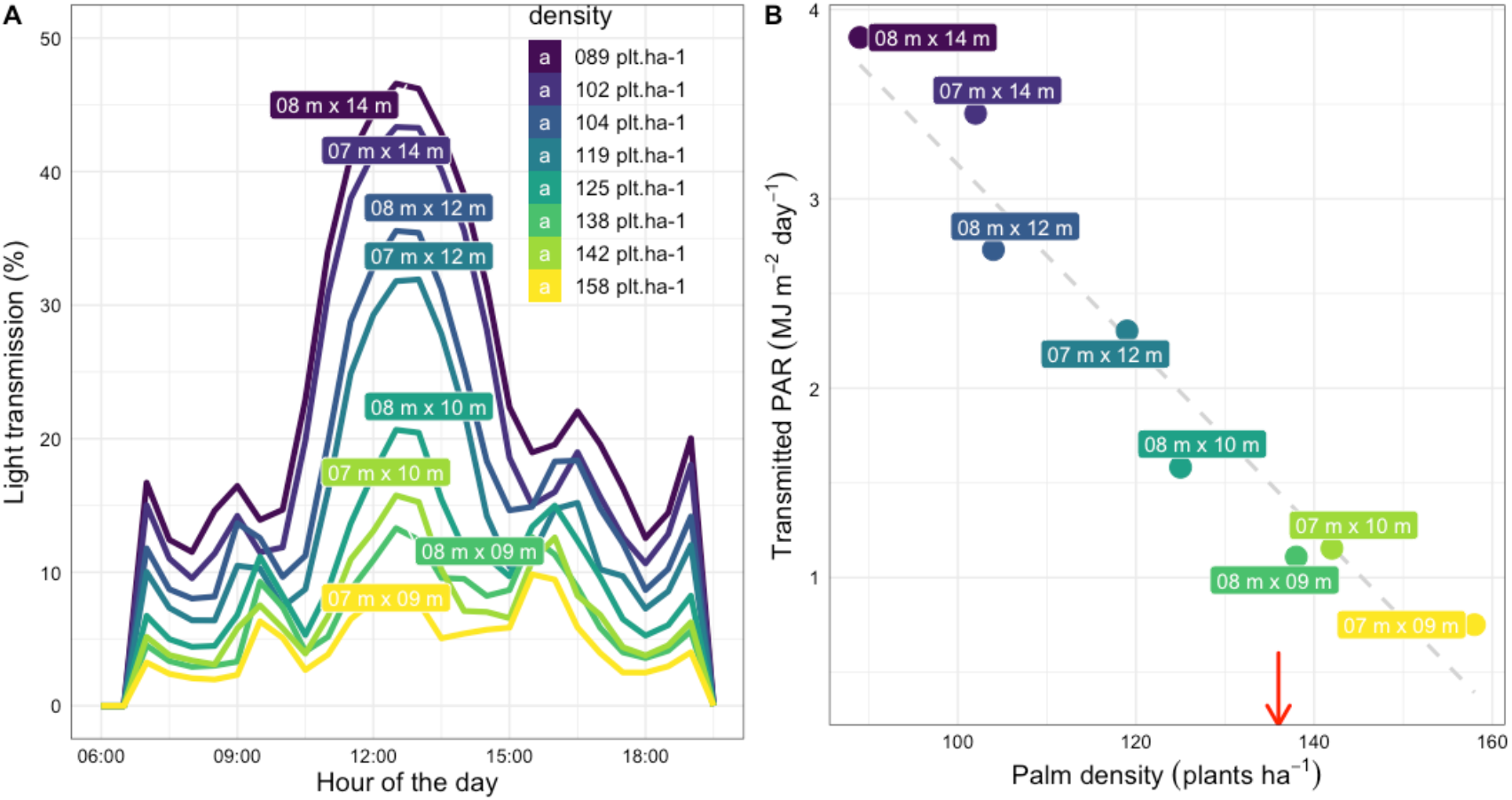
A) Average light transmission in the rice area over a day for the height simulated designs. B) Average daily photosynthetic active radiation available for rice between palm rows. The red arrow represents the conventional density (136 plants ha^-1^) and the dotted line the average loss of transmitted light with increasing density.

In one hand, the distance between rows strongly influenced the intensity of light around midday (Fig 4A), distances lower than 12m sharply limiting the maximum light transmission below 20%. In the other hand, distances over 12m decreased oil palm density below 110 plants ha^-1^. The 7 m x 12 m design exhibited the best compromise to ensure a sufficient palm density and limiting shading between rows. Moreover, this density of 119 plant ha^-1^ is close to the conventional densities of 136-143 plants ha^-1^. The 7 m x 12 m design was thus chosen as the reference to mimic in controlled conditions rice-palm agroforestry light, i.e the 60% of light attenuation over the day (Fig 1).

### Changes in plant morphology are mainly explained by daily light quantity while plant phenology is also impacted by light fluctuation

Under the shading treatments two varieties showed drastic increased in leaf length and etiolation symptoms leading to severe lodge, which prevented conducting them up to harvest: varieties V1 and V2 were thus removed from the three growth chambers 55 days after sowing. This first result pointed out how morphological responses to shade could introduce additional yield-reducing factor as lodging, and can be a first criterion to select adapted varieties. The variety V6 did not reach production maturity within the experimental time window, and the variety V4 was sensitive to the photoperiod chosen for the experiment, thus preventing plants to flower. Consequently, the following results will focus on the four other varieties on which phenotypic monitoring was performed from sowing to harvest.

Reduction in daily light quantity significantly affected the dynamic of leaf and tiller emission (Table 3 and Fig 5A & B). The phyllochron on the main stem increased under shade, with variations in the total number of leaves emitted depending on treatments and varieties (Table 4). During the first month after sowing, the number of leaves on the main stem was higher under the control treatment than under shade, but at maturity, plants under shade produced more leaves on the main stem for the *japonica* varieties (Fig 5A). The *indica* variety V3) emitted more leaves on the main stem under the control treatment than in the S treatment whereas the japonica V5 presented an opposite behavior with an increased number of leaves both shading treatments (Table 4). A significant effect of light fluctuation was revealed on phyllochron and the total number of leaves on the main stem, with important variations among varieties (Table 4). Regarding tiller emission, reduction in DLI induced significantly less tillers. DLF modulated the total number of tillers emitted, with more tillers in S-AF than in S, with significant differences for V5 and V8. Leaves length and width on the main stem sharply increased under both shading treatment (Fig 5C), and these changes in leaf dimensions were attributed to DLI and not to DLF (Table 3). The flag leaf shape (LW ratio) changed significantly for variety V3 with a higher length to width ratio under shade, while the other three varieties maintained a comparable shape and even tended to reduce the ratio by contrast (Table 4).

**Figure 5:**
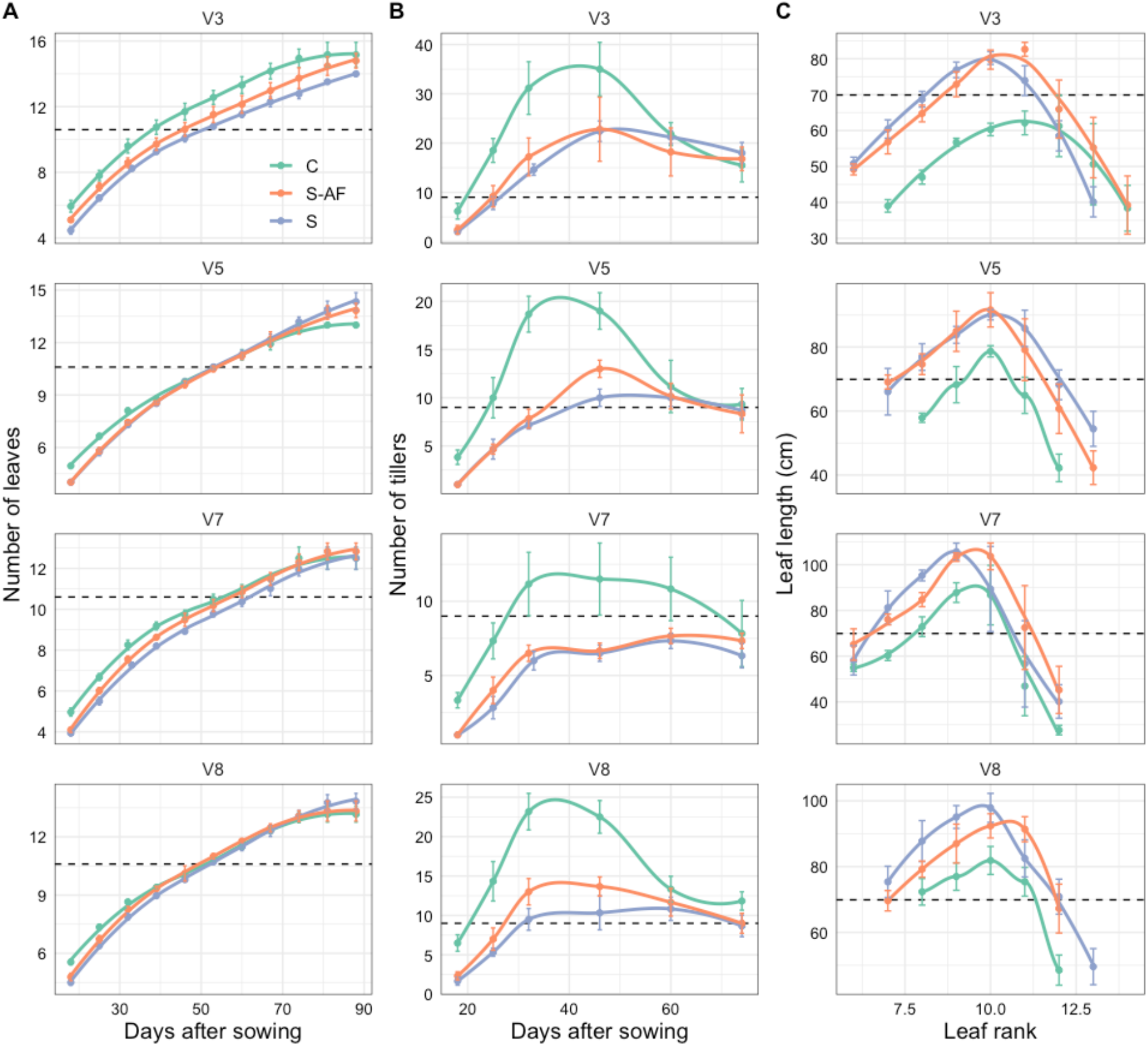
Dynamic of leaf emission according to varieties and treatments (A), dynamic of tillers emission according to varieties and treatments, (B) and length of leaves on the main stem (C). Smooth lines and points represent average values and vertical lines standard deviations. Dotted lines indicates the median value (taking all treatments and varieties together) for better visualize the difference between varieties.

**Table 4:** Phenotypic variations of traits with varieties and treatments. Numbers indicate mean value and letters represent significant difference between treatments for each variety assessed by Tuckey’s test (P<0.05)

|  |  |  |  |  |  |  |  |  |  |  |  |  |
| --- | --- | --- | --- | --- | --- | --- | --- | --- | --- | --- | --- | --- |
| 318<br>319 | Table 4: Phenotypic variations of traits with varieties and treatments. Numbers indicate mean value and letters represent significant difference between treatments for each variety assessed by Tuckey's test (P<0.05) |  |  |  |  |  |  |  |  |  |  |  |
|  | V3 |  |  | V5 |  |  | V7 |  |  | V8 |  |  |
|  | C | S | S-AF | C | S | S-AF | C | S | S-AF | C | S | S-AF |
| Yield | 34.2a | 28.7ab | 19.9b | 38.7a | 25.8b | 31.2b | 18.5a | 14.1a | 12.9a | 33.4a | 28.5ab | 23b |
| Nb_Pan | 14.8a | 13.0a | 12.8a | 9.33a | 7.33a | 7.67a | 6.5a | 5.17a | 4.83a | 10a | 7.67b | 8.33ab |
| Nb_Kern | 92.1a | 90.9a | 68a | 143a | 138a | 143a | 115a | 116a | 111a | 97.7b | **120a** | **89.1b** |
| TKW | 27.9a | 26.6a | 28.5a | 30.7a | **27.2b** | **30.7a** | 27.1a | 24.4b | 26ab | 36a | 32.8b | 32.2b |
| SF | 85.9a | **80.4a** | **59.5b** | 89.5a | 86.4a | 85a | 85.8ab | **94.3a** | **76.1b** | 94.2a | 92.6a | 90.2a |
| HI | 35.2a | **37.7a** | **21.3b** | 46.3a | **41.2c** | **43.4b** | 26a | 25.3a | 23a | 35.7b | **38.1a** | **33.8b** |
| DaysFL | 77.5b | 88a | 86.2a | 76.3b | 85.7a | 83.3a | 74b | 81a | 81a | 76.3b | 82.2a | 78.7ab |
| phylochron | 5.11c | **6.28a** | **5.75b** | 5.87a | 5.98a | 6.03a | 5.93b | 6.49a | 6.32a | 5.8a | 5.94a | 5.9a |
| Nb_Lea | 15.2a | **14b** | **15a** | 13b | 14.3a | 13.8a | 12.5a | 12.5a | 12.8a | 13.2a | 13.8a | 13.3a |
| Nb_Til | 35a | 22.6b | 25.2b | 19.3a | **10.5c** | **13b** | 11.5a | 7.33b | 7.67b | 23.7a | **11c** | **14.2b** |
| CPN | 0.32a | 0.23b | 0.21b | 0.24a | **0.12c** | **0.16b** | 0.19a | 0.14b | 0.14b | 0.19a | 0.14b | 0.14b |
| PH | 88.8a | 82.3a | 97.2a | 113c | **121b** | **128a** | 137a | 141a | 148a | 110b | 123a | 117ab |
| VDW | 60a | **44.8b** | **68.8a** | 43a | 35.6a | 38.9a | 51.9a | 40.4a | 42.3a | 58.2a | 45.1b | 44.3b |
| PLA | 2316b | 3279ab | 3348a | 2173a | 2223a | 2487a | 1615b | 2117ab | 2515a | 2063a | 2552a | 2630a |
| SLA | 209b | 252a | 263a | 177b | 218a | 220a | 158b | 229a | 216a | 165a | 210a | 215a |
| FL Length | 31.4b | 40.3a | 40a | 38.1b | 47.2a | 40.9ab | 32.9b | 41.6a | 43.7a | 46.3b | 54.6a | 56.7a |
| FL Width | 1.33b | 1.51a | 1.46a | 1.39b | 1.79a | 1.76a | 1.69b | 2.15a | 2.12a | 1.54b | 1.94a | 1.83a |
| FL LWratio | 23.7b | 26.7ab | 27.2a | 27.6a | 26.5a | 23.3a | 19.4a | 19.2a | 20.6a | 29.9a | 28a | 31a |
| FL Area | 30.4b | 45.2a | 45.4a | 40.5b | 60.8a | 50.7ab | 37.8b | 60.4a | 59.5a | 48b | 75.2a | 73.4a |
| FL Incl | 168a | 165a | 166a | 164a | 161a | 162a | 93.8ab | **107a** | **63.2b** | 160a | **128b** | **146a** |
| FL StoDen | 748a | 698a | 671a | 641a | 623a | 649a | 594a | 531ab | 516b | 583a | 512b | 544ab |
| SGA | 70.8b | 150a | 136a | 3.66c | **176a** | **68.9b** | 68b | 344a | 305a | 16.2c | **165a** | **119b** |
| SPAD | 31.9a | 34.3a | 37.1a | 35.4b | **43.1a** | **36.7b** | 41.6a | 41.2a | 35.3a | 36.2a | 36.1a | 36.2a |
| Lcp | 55.6a | **67.1a** | **33.1b** | 34.2a | 22.3a | 31.2a | 39.6a | **29a** | **16.7b** | 30.8a | 32.4a | 35.7a |
| Asat | 16.6a | 15.6a | 20.3a | 13.7b | **19.7a** | **11.7b** | 17.2a | 15.1ab | 11.6b | 18.1a | 17.6a | 15.4a |
| Aqe | 0.0021b | **0.0018b** | **0.0028a** | 0.0027ab | **0.0033a** | **0.0023b** | 0.0029b | **0.0028b** | **0.0042a** | 0.0031a | 0.0029a | 0.0027a |
| Rd | 0.50a | 0.42a | 0.81a | 0.57ab | **0.67a** | **0.25b** | 0.75a | 0.64a | 0.34a | 0.9a | 0.85a | 0.63a |
| ETRm | 154a | 142a | 166a | 153ab | **173a** | **136b** | 163a | 149ab | 121b | 168a | 159a | 152a |
| alpha | 0.29ab | **0.27b** | **0.32a** | 0.32a | **0.33a** | **0.3b** | 0.33ab | **0.32b** | **0.34a** | 0.33a | 0.32a | 0.31a |
| lk | 522a | 547a | 516a | 476a | 526a | 458a | 491a | **469a** | **356b** | 505a | 500a | 490a |
| M | 6.71a | 6.67a | 6.68a | 6.22a | 6.55a | 6.33a | 4.73a | 4.99a | 4.67a | 6.79a | 6.57a | 6.65a |
| m | 2.35a | 2.25a | 2.25a | 2.43a | 2.55a | 2.44a | 2.70a | 2.67a | 2.69a | 2.60a | 2.54a | 2.55a |
| Star % | 77.5a | 77.2a | 78.4a | 84.5a | 72b | 79.2ab | 84.6a | 77.6a | 75.4a | 76a | 68.6a | 77.8a |
| Amy % | 23.5a | 25a | 23.7a | 14.2a | 14.5a | 14.8a | 21a | 21.7a | 21.6a | 12.8a | 12.6a | 10.6a |
| ε % | 10.3a | 9.60a | 9.78a | 8.45b | **10.7a** | **8.38b** | 11.7a | **9.54c** | **10.3b** | 8.66ab | **8.82a** | **8.29b** |

Interestingly, stomatal density on flag leaf significantly decreased with DLI, but only for the two varieties V7 and V8, while the two others varieties V3 and V5 kept a comparable stomatal density while increasing flag leaf area. At the plant level, the decrease in DLI involved taller and less erected plants (lower CPN), mainly due to longer, wider and thinner leaves as well as longer internodes (Fig S4).

The total plant leaf area increased under shading treatments without marked difference between S and S-AF. The SLA increased in both shading treatments without marked difference du to DLF, traducing the effect of light quantity alone in leaf elongation and widening. The effect of DLF was only important for the total vegetative dry matter (VDW) at harvesting, partly because of the contrasted behavior of V3 between S and S-AF (Table 4).

### Light fluctuation impacted leaf senescence and grain filling, and involved high variation in photosynthetic capacities among varieties

At the plant scale, the most marked difference between treatments was the stay green area at physiological maturity (Table 3 & Table 4). We observe a gradient in leaf senescence with light fluctuation, plants from the control treatment presenting higher senescence than plants under shade that stayed greener. Increasing DLF significantly increased senescence (low SGA), suggesting that resource reallocation may depend on irradiance intensity.

Plant yield was significantly higher in the control treatment, and no major differences were stressed between the two shading treatments (Tables 3 & 4). Lower yield under shade was mainly explained by the reduction in the number of panicles per plant, but the establishment of grain yield varied with DLF. Indeed, the comparison of production between S and S-AF revealed that under higher light irradiance (S-AF), spikelet fertility was reduced, thus limiting the number of kernels per panicle. Nevertheless, yield was maintained under S-AF trough higher TKW (bigger grains) (Table 3), suggesting that grain filling was enhanced with irradiance intensity. The varieties V3, V5 and V8 produced more than the variety V7 in all the treatments. Interactions between varieties and shade treatments was stressed, the variety V5 producing significantly more than V3 and V8 under S-AF conditions while no significant difference was stressed between these three varieties in the two other treatments (Fig 6).

**Figure 6:**
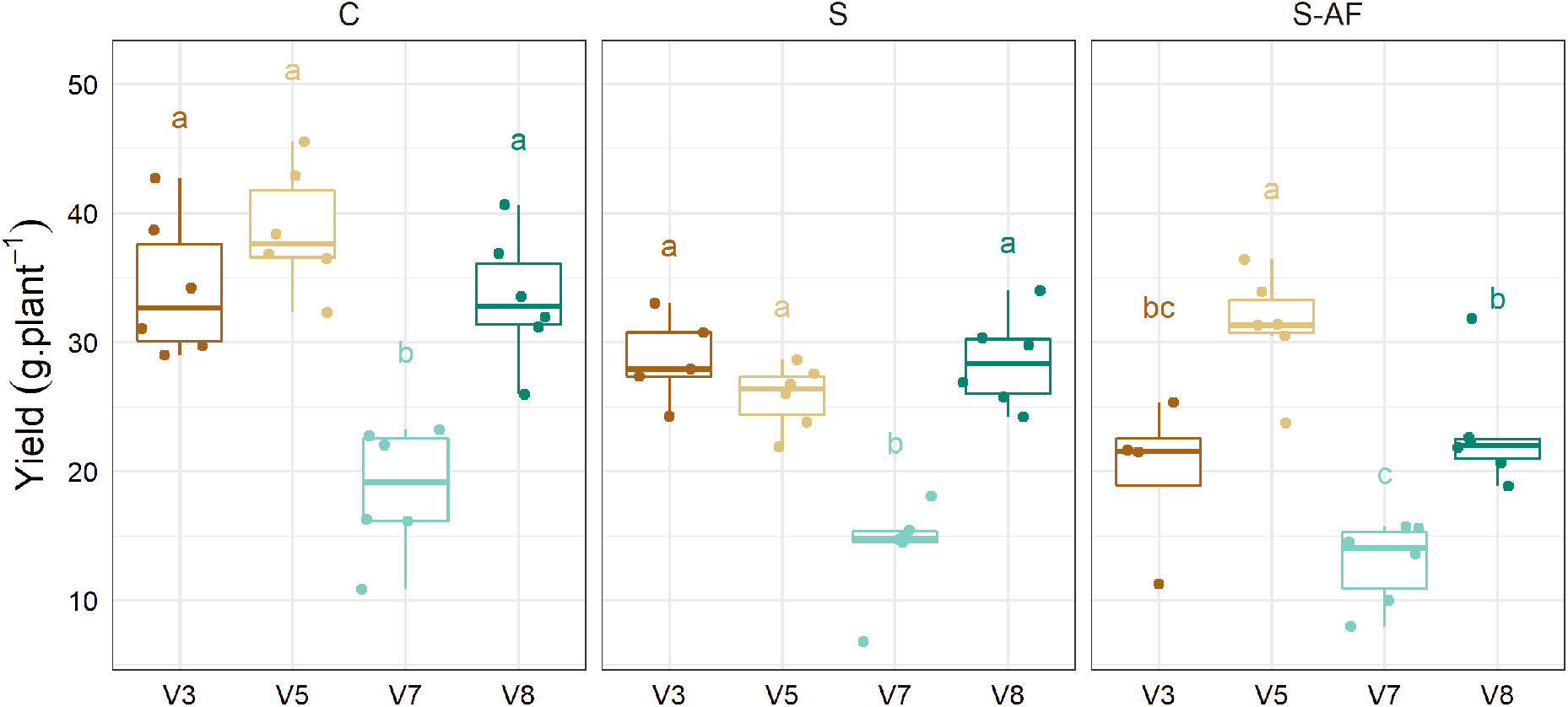
Plant yield for the four varieties studied in the three light treatments. Letters indicate significant differences between varieties in each treatment (Tuckey s test P<0.05).

The study of flag leaf CO^2^ assimilation revealed high interaction effects (V x DLF) for nearly all photosynthesis parameters (Table 3). Light-responses curves exhibited contrasted behavior of varieties, with more varietal variability under the S-AF treatment than in the two other treatments (Fig 7). The variety with maximal photosynthesis was different in each treatment, revealing the high variability in photosynthetic processes with environment and the potential weak heritability of photosynthetic traits. No clear relationship could be established between photosynthetic capacity and other morphological or yield-related variables, highlighting the complexity and multifactorial processes involved in plant growth and production. At the grain scale, the morphology of grain (length and width) was not significantly affected by the light treatments (Table 4). Complementary ANCOVA confirmed that DLI and DLF has no significant effect on grain morphology (Table 3) contrarily to the factor variety, suggesting that kernel length and width are heritable traits.

**Figure 7:**
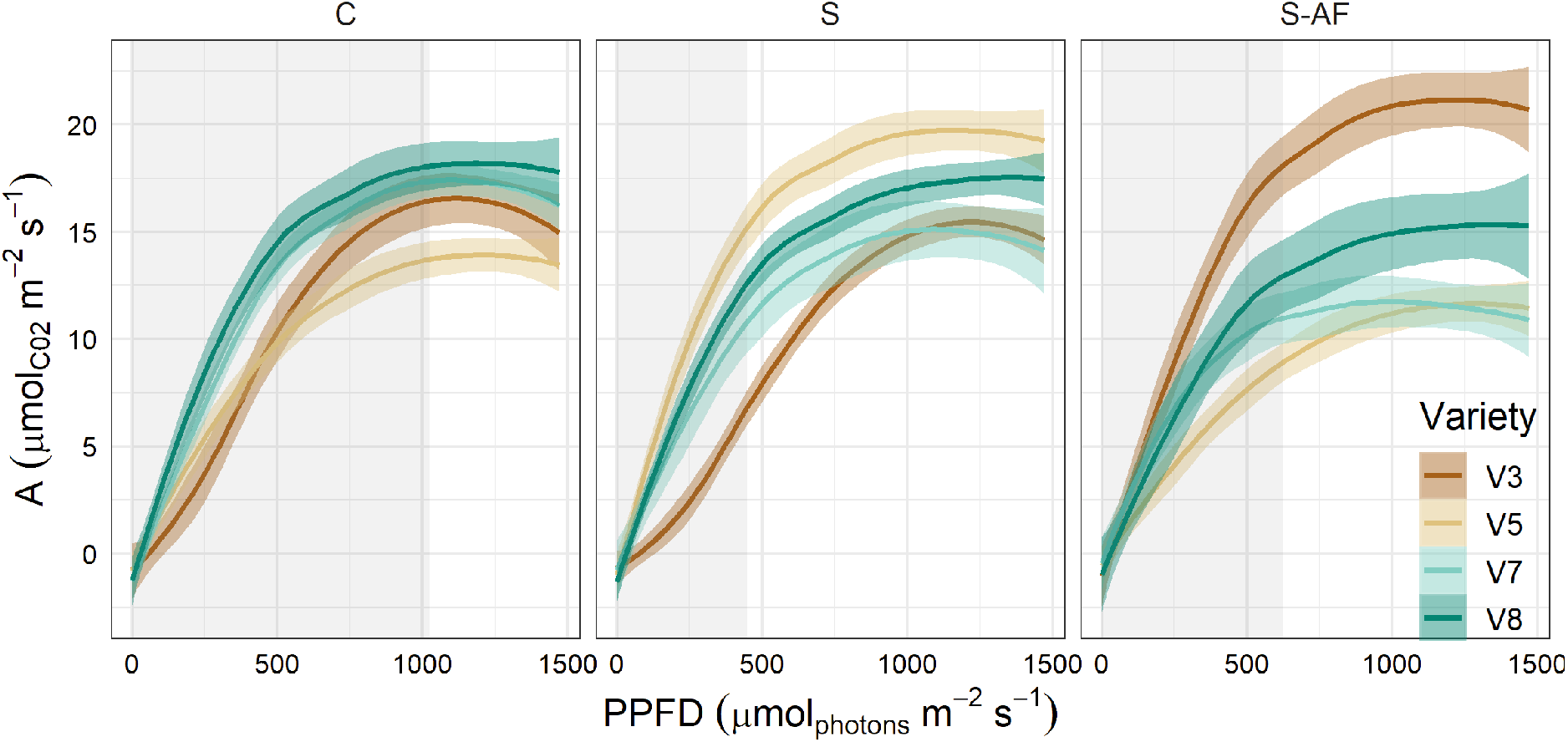
Response curves of carbon assimilation with irradiance according to treatments and varieties. Lines indicate smooth curves adjusted per variety and per treatment with 95% confidence interval. Grey areas represent the light fluctuation domain provided in each treatment.

Both amylose and starch content were revealed being also significantly affected among varieties. The variation of the composition of starch was only observed among V5 under low light, where the DLI treatment induced a significant reduction of the amylose content as compared to the control one. Varieties V3, V5, V8 exhibited significant variation in their porosity. The kernel porosity was significantly affected by varieties, by treatments and their interactions (Table 4). Contrary to V5 and V8 varieties, V7 presented a significantly lower level of air space for the kernel grown under S than that of S-AF environment.

### Relative changes among varieties are maintained between the light treatments

Three principal component analysis (PCA) were performed, one for each treatment (Fig 8). For all PCA, more than 60 % of the phenotypic variance was explained by the two first components (PC1 & PC2), the contribution of PC3 to total variance being markedly lower than PC2. In the control treatment (Fig 8A), vegetative components (number of tillers, number of leaves) were the major contributors to PC1 while yield components (kernel weight, harvest index) contributed to PC2.

**Fig 8:**
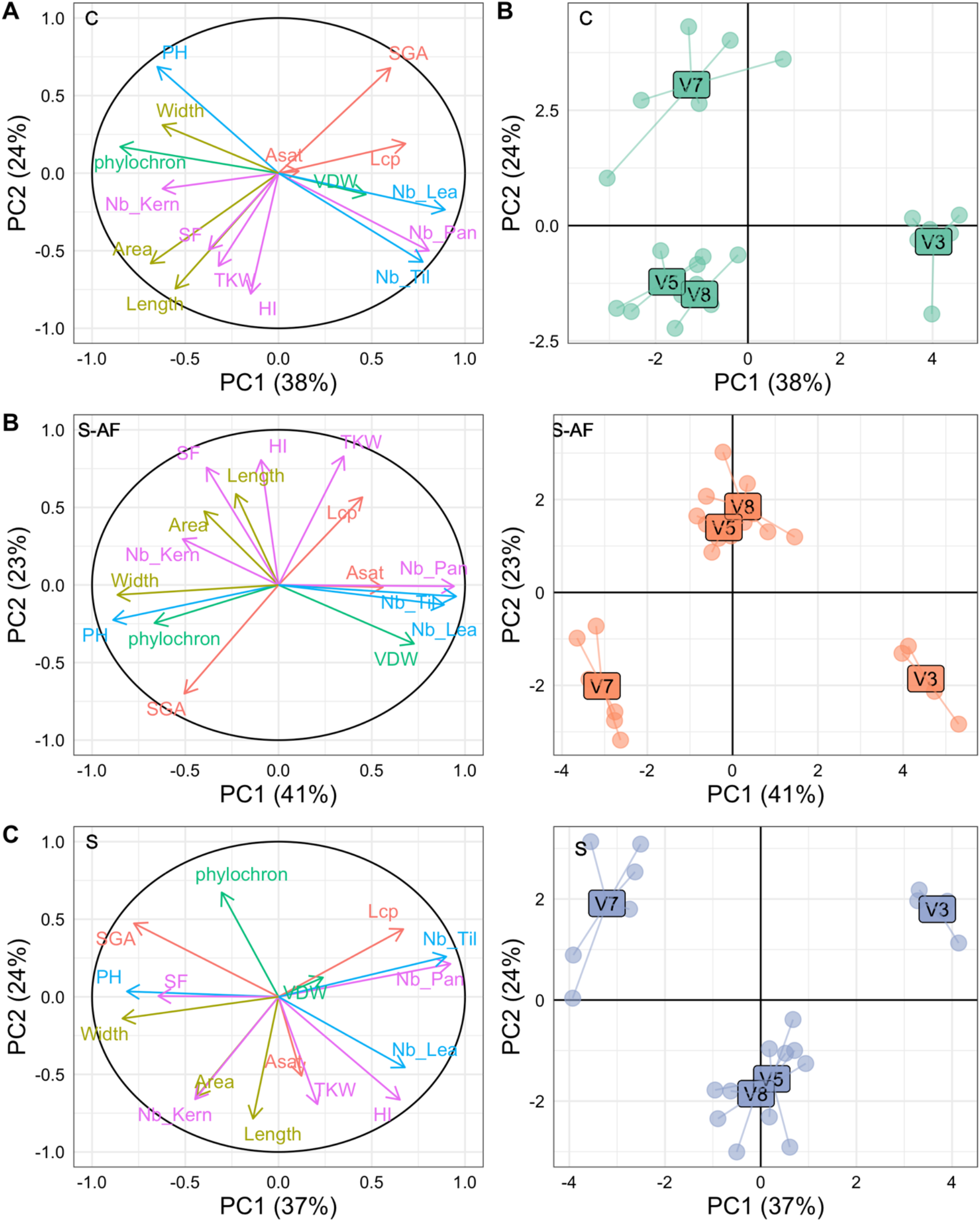
Principal component analyses performed on the three treatments (A: treatment C, B: treatment S-AF and C: treatment S). Graphics on the left represent the projection of traits on PCA axes and graphics on the right the projection on PCA of individuals grouped per varieties

Interestingly, flag leaf morphology (length and area) contributed more to PC2 than PC1, suggesting a closer link between flag leaf geometry and yield components than between the whole plant vegetative attributes and yield. PCA for S-AF (Fig 8B) resulted in similar interpretations, the vegetative components contributing to PC1 and yield components to PC2. Additionally, plant height and flag leaf width contributed markedly to PC1, which revealed the architectural variations triggered by lower light intensity. Physiological variables such as Lcp, Aqe and SGA were highly correlated to yield components. Plants with the highest yield presented low SGA (high senescence), low photosynthetic efficiency and high light compensation point. This result confirmed that under S-AF treatment, yield depended more on resource remobilization to the panicle rather than enhancement of the photosynthetic capacities of the flag leaf. The PCA performed on the treatment S (Fig 8C) indicated only small discrepancies with the PCA for the S-AF. The dichotomy between vegetative/architectural features and yield features was marked in both shading treatments. Variations in flag leaf width particularly discriminated individuals in shading treatments, and in contrast to the control treatment, a higher correlation was observed between flag leaf with and yield than between flag leaf length and yield (Fig S5). Projection of individuals on the three PCAs clearly split varieties, revealing that the studied phenotypic traits and their combination were specific to varieties. In the three treatments, projections of V5 and V8 were superposed, denoting the phenotypic similarities between those two varieties. Architectural features of V3 and V7 were significantly different since there projections were in opposition in respect to PC1, notably in the shading treatments. Finally, the similarity between varieties projections from the three PCAs may suggest that varietal responses to low light are likely to be heritable, and that varietal screening under full light can provide clue on varietal behavior under low light.

## Discussion

### Using functional-structural plant models to design innovative agroforestry systems

The development of agroforestry system often depends on the success of smallholders’ initiatives to test new farming practices. The adoption of oil palm agroforestry systems has been promoted among smallhoders through empirical practices conferring socio-economic and environmental benefits (Susanti *et al*. 2020). However, it is not straightforward to conduct specific studies to understand and optimize the agronomic performance of oil palm agroforestry systems, since such studies require years of investigation and are very costly. Intercropping is generally only associated in the first years after planting, in systems where the density of oil palm is optimized for growing the palm alone at maturity (Koussihouèdé *et al*. 2020). In this study, we considered that future food security and environmental concerns would require the development of sustainable oil palm agroforestry systems, with the possibility of growing food crop in association with mature oil palms.

In this context, we proposed a new approach, based on the assessment of light distribution in a mature oil palm agroforestry system. Using a quantitative trade-off between light resource for rice cultivation and oil palm density to optimize oil production, the modelling approach proposed in this paper showed how FSPM could help designing oil palm-based intercropping systems with scientific background. The proposed methodology could be adapted to many systems that requires time-consuming and expensive trials. However, it is worthy to note that this approach was particularly suitable for oil palm-based systems due to the low architectural plasticity of the oil palm, and its low impact on light interception efficiency at the plot level (Perez *et al*. 2022). The low plasticity of oil palm reinforces its suitability for association with other crops. In this study we did not consider the effect of planting density on oil palm architecture and subsequently light interception, but it is likely that the same approach with perennial plants with complex branching patterns and highly plastic to the radiative environment would require additional modeling inputs to derive reliable light partitioning predictions. The number of designs tested in the presented simulation study was limited to a few numbers, but further simulations could be performed to mathematically optimize light partitioning and oil palm density, including new patterns different from the row planting proposed here.

Studies focusing on the effect of light attenuation are often performed in outdoor conditions using shading net (Liu *et al*. 2014; Chen *et al*. 2019; Jia *et al*. 2021), and more recently in growth chambers (Yang *et al*. 2018; Zhang *et al*. 2020). Highly controlled growth chambers are generally used to simulate future climate conditions (i.e rising CO2 or temperature), but can now be used to simulate change in light regime (Vialet-Chabrand *et al*. 2017). The originality of our study was to use modeling outputs to set up light treatments in controlled conditions. Conversely to outdoor conditions where the dynamic of light cannot be controlled, here we were able to investigate the impact of both light quantity and the dynamic of light intensity. Such experimental device aimed at mimicking the light regime of an agroforestry system not yet implemented, taking into account the reduction of light and its variation over the day. However, the major drawback of our experiment was the inability to modulate light quality (red/far red ratio) as it would be in the shade of oil palm, which can affect growth and development of the understorey crops (Chen *et al*. 2019; Zhang *et al*. 2020).

### Toward the selection of rice varieties adapted to agroforestry light conditions

Three characteristics of light are modulated under agroforestry systems: the quantity of light, the fluctuation of light, and the quality of light. The inexistence of real conditions to screen the tolerance of rice varieties to the agroforestry system of interest involved setting up a trial in controlled conditions mimicking these conditions. Our controlled conditions prevented us from modulating light quality, and more precisely the red to far-red ratio, which is known to be a light signal triggering a suite of physiological and morphological responses called the shade avoidance syndrome (Franklin 2008; Ballaré & Pierik 2017). Zhen and Bugbee (Zhen & Bugbee 2020) recently stressed the importance of far-red photons on plant photosynthesis, suggesting the need to integrate a larger range of wavelength in studies dedicated to investigate responses to light. Other studies already demonstrated the genetic inheritance of cereal crops in responses light quality (Wille *et al*. 2017), but only few investigated the interaction between light quantity and quality (Yang *et al*. 2018). Further dedicated experiments integrating response to light quality, in combination with light quantity and light fluctuation would thus be of great interest to complement the result found in the present study and better understand responses to agroforestry light regime. However, in the intercropping design prospected here, change in light quality would appear only a few hours during the day. This part time change in light quality raises new question about how long the light signal must last to trigger the shade avoidance syndrome (Li *et al*. 2021).

In our experiments, we focused on the effect of light integral (DLI) and light fluctuation (DLF) on rice growth and production. DLI represented an index of resource availability, whereas DLF was considered as a light signal. Our results confirmed that the reduction in the quantity of light on rice prolonged the period of growth, increased plant height and leaf area, decreased the number of tillers and total plant dry mater (Liu *et al*. 2014) and subsequently reduced grain yield (Wang et al. 2015b; Chen et al. 2019). Our result clearly showed the changes in resource investment under low light: plants produced fewer tillers but increase the number and the size of leaves in the main stem. Shading treatments triggered leaf elongation and widening, conferring a higher leaf area than under high light intensity. In parallel, leaves were thinner under shade, traducing a lower capacity to accumulate nitrogen (Murchie et al. 2005). The adjustment in leaf morphology were consistent with other studies in rice (Liu *et al*. 2014) or others crops (Arenas-Corraliza *et al*. 2019; Poorter *et al*. 2019), notably the increase of SLA, but showed some discrepancies with respect to leaf widening. Contrarily to the present study and the review by Liu (Liu *et al*. 2014), reductions in leaf width under shade were observed for rice (Lafarge *et al*. 2010) and for maize (Lacube *et al*. 2017). These discrepencies could be explained by the duration and the intensity of shade, and raises question about the existence of light signal thresholds that trigger morphological adjustments.

Light quantity was the main factor driving morphological and architectural changes, while light fluctuations only appeared to explain variations in yield components and phenology (Table 3). This study highlighted the importance of light fluctuation in the grain filling process and resource reallocation. The stay green area was particularly affected by the intensity of light experienced by the plant, revealing that fluctuating light matters to trigger some process linked to resource reallocation and subsequently plant yield. Here we demonstrated that light fluctuation is part of the light signaling pathway that regulates leaf senescence (Woo *et al*. 2019), and that stay green can be an informative trait for the selection of varieties adapted to agroforestry systems. The reaction between light intensity and reallocation patterns have already been observed in wheat with vegetative dry matter redistribution into grains (Li *et al*. 2010), and with nitrogen reallocation in the leaves to optimize photosynthetic performance exposure (Li *et al*. 2021).

Among others, it has been established that low light intensity under shading nets could affect the grain quality traits, including the starch content its and composition (Li *et al*. 2006; Wang *et al*. 2013; Liu *et al*. 2014) and grain appearance. Our results confirmed that light quantity affected starch content in the grain, without altering the fraction of amylose. Wang et al. (2013) earlier reported some significant differences in total starch content and amylose content of milled rice flour, and concluded to the contribution of heredity, environment and their interaction for indica hybrid rice varieties under shading. The decrease in the amount of starch and amylose was also claimed by Li et al. (2006) in two grain varieties indica and japonica under shading treatments with nets (50% transmittance rate). The presence of chalk, which is a key criterion for visual assessment of grain marketability, can also be affected by shading environmental-related stress (Liu *et al*. 2014; Patindol *et al*. 2015). High porosity is usually related to the presence of chalk (Singh *et al*. 2003), and characterized by the presence of disorganized cellular structure (Lisle *et al*. 2000) leading to a greater cell and granular swelling, detrimental to grain eating quality. The four studied varieties showed contrasted grain porosity in responses to light quantity and fluctuation. The indica variety tended to present a lower porosity under both shading treatments while the three other varieties showed either higher or lower porosity than the control treatment. In this study, the significant loss in yield under shading treatments did not result in a significant decrease in milling quality as it observed in previous studies (Liu *et al*. 2014). Further research would be needed to investigate the qualitative response of rice plants to fluctuating radiations and shading, whose tolerance might be genotype-dependent. It is likely to favor the selection of suitable cultivars to meet the expectations of end-users, under constrained agroforestry growing environment.

The photosynthetic capacity measured via light response curve highlighted that light quantity and fluctuation triggered different responses among the four studied varieties. The high interaction between varieties and shading treatments was already stressed in previous studies (Wang *et al*. 2015b; Pan *et al*. 2016), revealing the difficulty to select adapted varieties trough photosynthetic measurements. The physiological measurements performed did not help retrieve the difference in production among varieties, suggesting that yield is the consequence of multifactorial processes that can compensate variations in photosynthetic capacity. These specific responses to light intensity and fluctuation did not provide substantial benefits in term of production. Nevertheless, the absence of evolution of the morphological kernel traits under shading stressed that kernel shape is highly heritable and irresponsive to change in light intensity and fluctuation. It is likely that leaf scale measurements did not reflect an average value of the whole canopy photosynthesis. Besides, photosynthesis measurements are time consuming, preventing the efficient screening of varieties for breeding purpose.

Even if marked interactions were observed in physiological traits among varieties and treatments, the yield ranking between the four varieties was maintained in the three treatments. From an experimental and phenotyping perspective, this result tends to support that artificial light depletion using shade net could be sufficient to screen varieties of interest for agroforestry systems not yet implemented. Our result also confirmed previous findings revealing that productive materials in full sun conditions are also good producers in low radiation (Bueno & Lafarge 2017). The very specific behavior and low production (with potential low cooking and eating quality) of V7 together with the comparable production of the three other varieties made the identification of traits of interest difficult, without implementing reliable and discriminating descriptors. Indeed, the contrasted traits values between the two groups of individuals, formed by V7 and the three other varieties, could bias trait correlations. However, the variety V5 showed a significant higher production than the other varieties in the S-AF treatment, without exhibiting a chalky endosperm appearance potentially detrimental to its grain marketability. This variety maintained the highest harvest index among all the varieties and treatments, with a limited expansion of vegetative biomass in response to shade and a high leaf senescence under shade. These characteristics are consistent with the shade tolerant responses optimizing the trade-off between the elongation of vegetative parts and yield (Wille *et al*. 2017), and pave the way for selecting varieties dedicated to agroforestry systems. Further research would be needed to enlarge the diversity of the genotypes to screen and clearly identify if specific traits are better adapted to agroforestry systems. Other perspectives studies would also need to integrate other factors than the light resource, including a more global analysis with the environmental, social and economic aspects of the agroforestry system proposed (Rodenburg *et al*. 2022).

## Acknowledgments

The authors would like to thank the Montpellier GAMET Biological Resource Centre (Centre de Ressource Centre de Ressources Biologiques des “Graines Adaptées aux conditions Méditerranéennes et Tropicales », ARCAD Montpellier – UMR AGAP Institute de Montpellier) for providing the germplasm. The authors would like to thank Clémence Foray and Matthieu Dejean for the grain image acquisition, and for morphological script using open source ImageJ program, Romain Domingo for assistance in grain quality analyses. This research was supported by the General Directorate for Research & Strategy of CIRAD with incentive action “Scientific Creativity & Innovation” (CreSi) to both the Agroforice (RP), and the voxOmic projects (OG). RP and RV have been supported by the MaCS4Plants CIRAD network, initiated from the AGAP Institute and AMAP joint research units.

