## supplementary material for "Combining modelling and experimental approaches to assess the feasibility of developing rice-oil palm agroforestry system"

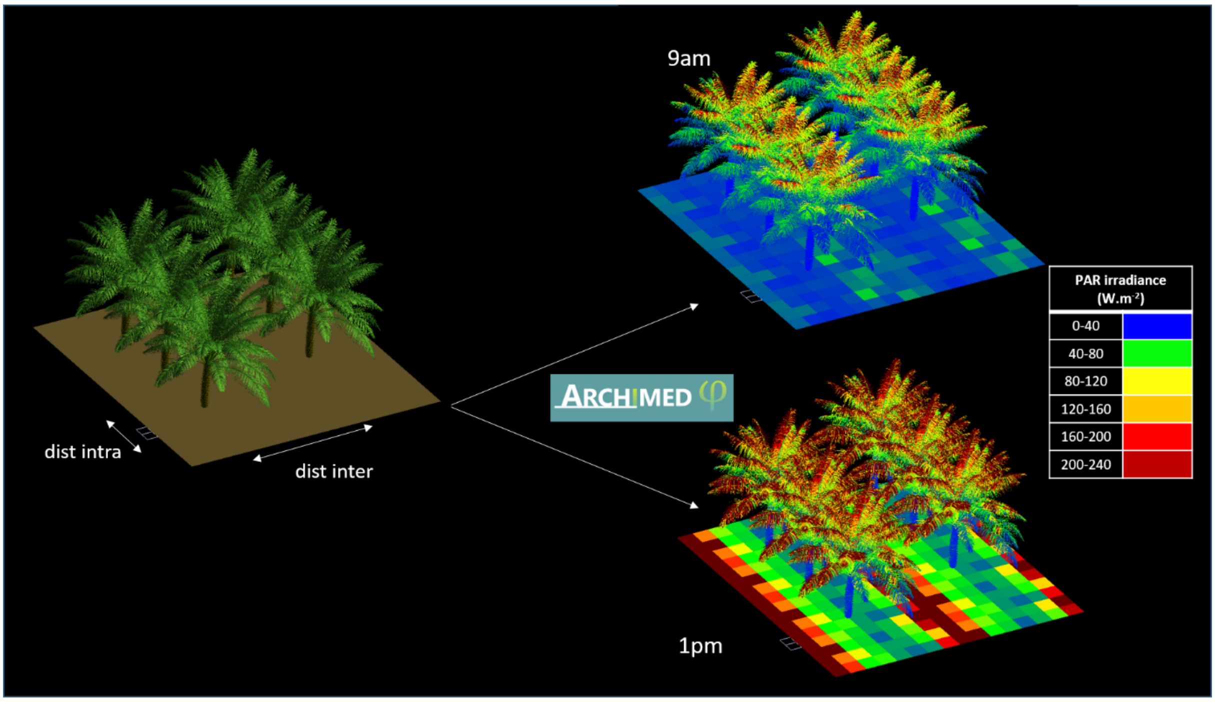


*Fig S1: Simulation of light interception on 3D mock ups of oil palm at two hours of the day (9am and 1pm). Each ground square integrates the light received over one hour on one square meter. Light interception is estimated on 3D scenes infinitely replicated (toricity) to avoid border effect.*

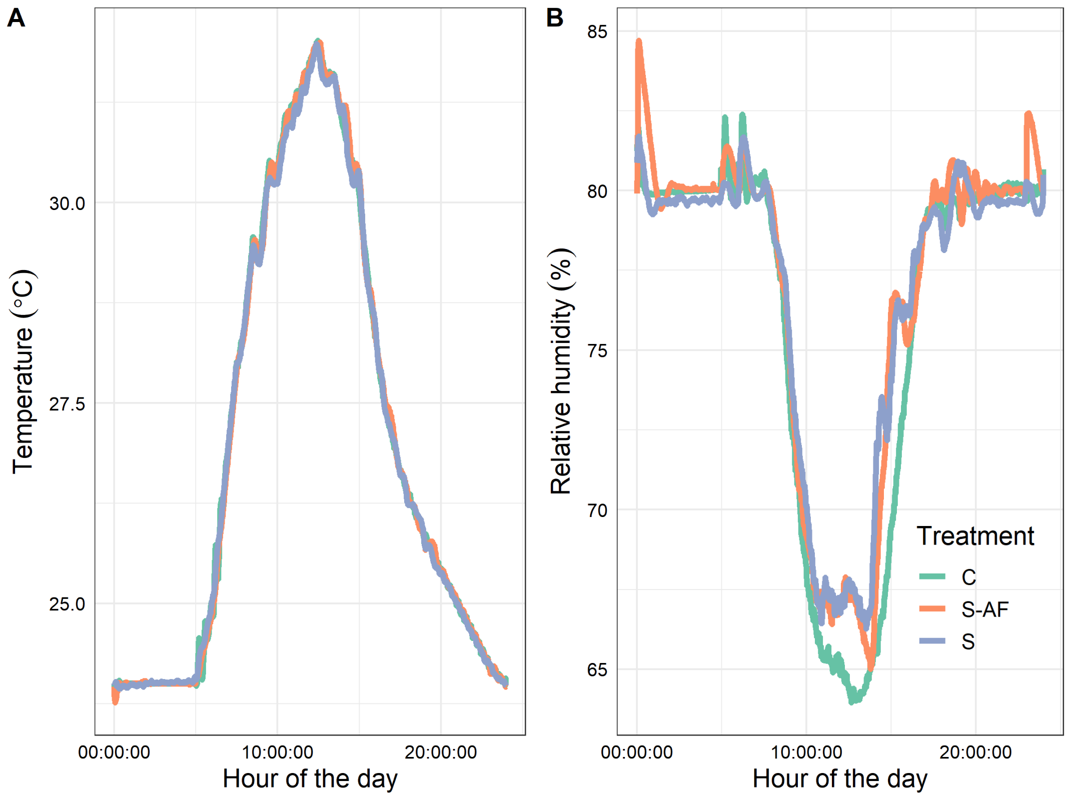


*Figure S2: Mean timecourse of temperature (A) and relative humidity (B) measured in the three phytotrons during the whole cycle of plant growth (C: control, S-AF: agroforestry-like shade, S: constant shade).*

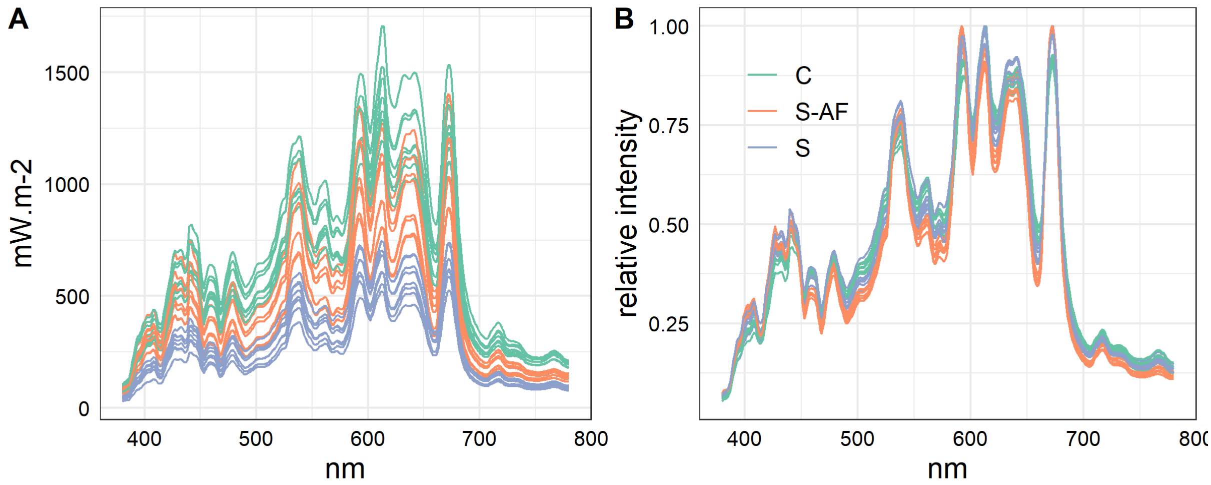


*Figure S3: Measurement of light spectrum in the three phytotrons with absolute light intensity (A) and relatively to the maximal intensity along the spectrum (B). Lines represent different positions in the phytotrons, measurements were performed around daily maximum light intensity (11am).*

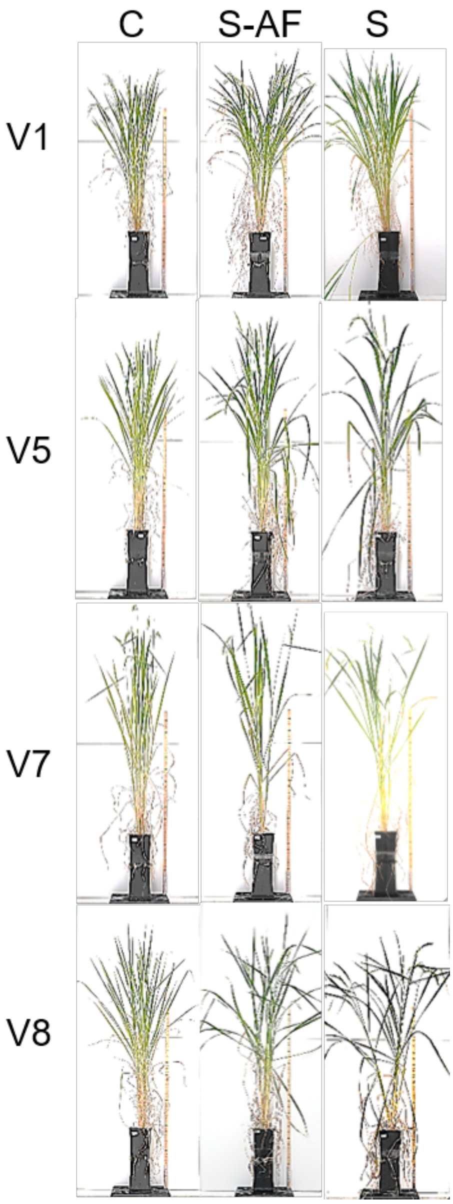


*Figure S4: Effect of light treatments on plant architecture depending on varieties (pictures were taken before harvesting).*

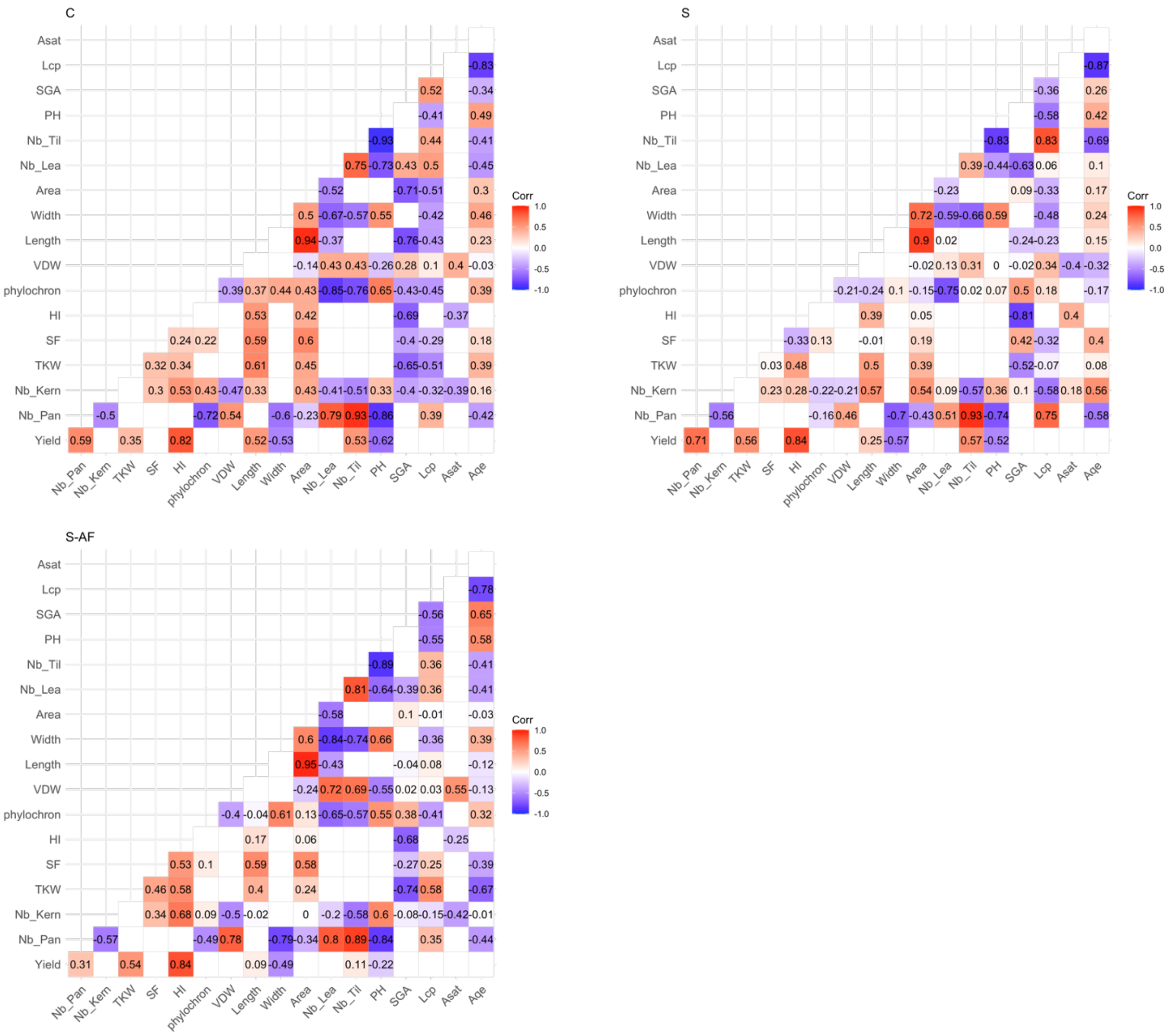


*Figure S5: Trait correlations in the three light treatments. Correlation coefficients are indicated with values and colors (blue -1, red +1)*
